# High-content image-based drug screen identifies a clinical compound against cell transmission of adenovirus

**DOI:** 10.1101/2020.03.24.006346

**Authors:** Fanny Georgi, Fabien Kuttler, Luca Murer, Vardan Andriasyan, Robert Witte, Artur Yakimovich, Gerardo Turcatti, Urs F Greber

## Abstract

Human adenoviruses (HAdVs) are fatal to immuno-suppressed people, but no effective anti-HAdV therapy is available. Here, we present a novel image-based high-throughput screening (HTS) platform, which scores the full viral replication cycle from virus entry to dissemination of progeny. We analysed 1,280 small molecular weight compounds of the Prestwick Chemical Library (PCL) for interference with HAdV-C2 infection in a quadruplicate, blinded format, and included robust image analyses, and hit filtering. We present the entire set of the screening data including all the images, image analyses and data processing pipelines. The data are made available at the Image Data Repository (IDR) ^1^, accession number idr0081. Our screen identified Nelfinavir mesylate as an inhibitor of HAdV-C2 multi-round plaque formation, but not single round infection. Nelfinavir has been FDA-approved for anti-retroviral therapy in humans. Our results underscore the power of image-based full cycle infection assays in identifying viral inhibitors with clinical potential.

## Background & Summary

Human adenoviruses (HAdVs) affect the respiratory, urinary and gastrointestinal tracts and the eyes. They cause morbidity and mortality, especially to immuno-compromised patients ^2,3^ as indicated by a recent outbreak in the USA killing 12 children, or a recent case of meningoencephalitis in a middle-aged woman in the US ^4^. HAdVs have a high prevalence ^5–8^ and are broadly used as gene therapy and vaccination vectors as well as oncolytic viruses ^9–11^. The high seroprevalence of HAdV-C2 and C5 (species C, types 2 and 5) underlines that HAdV infections are asymptomatic in healthy individuals, but persist in mucosal lymphocytes, and thereby pose a risk for immunosuppressed patients undergoing stem cell transplantation ^12,13^. More than 100 HAdV genotypes are grouped into seven species based on hemagglutination assays and genome sequences ^14,15^. Types of the species A, F and G replicate in the gastrointestinal tract, B, C and E in the respiratory tract, and B and D in the conjunctiva of the eyes. Notably, species B members have a broad tropism, including kidney and hematopoietic cells ^7,13^.

HAdV has a double-stranded DNA genome of ~36 kbp tightly packaged into an icosahedral protein capsid of about 90 nm in diameter ^16,17^. HAdV-C2 and C5 enter cells by receptor-mediated endocytosis, shed minor capsid proteins, expose the membrane lytic protein, penetrate the endosomal membrane and are transported to the nuclear membrane, where they uncoat and release their genome to the nucleus ^18–21^. In the nucleus, the viral genome gives rise to the immediate early viral mRNA encoding the E1A protein which transactivates the subviral promoters, drives lytic infection and maintains genome persistence in presence of interferon ^22–24^. Proteolytically matured HAdV progeny is released upon rupture of the nuclear envelope and the plasma membrane ^25–27^.

Currently, there is no effective therapy available against HAdV disease. The standard of care is the nucleoside analogue Cidofovir, with poor clinical efficacy ^7,28^. The problem is exacerbated by the shortage of a suitable small animal model for HAdV disease, although Syrian Hamsters are susceptible to HAdV-C infection and give rise to viral progeny ^29^. Here, we developed an image-based procedure to identify novel inhibitors of HAdV infection in cell culture. We used the commercially available Prestwick Chemical Library (PCL) comprising 1,280 off-patent mostly FDA-approved small molecules (listed in Supplementary Table 1). The PCL comprises compounds against diseases including infection and cancer ^30–32^.

Here, we performed a phenotypic screen against HAdV-C2 infection employing automated fluorescence microscopy and image-based scoring of the progression of multi-round infections using the Plaque2.0 software ^33^ (Figure 1a and b). The screen was performed in 384-well plates (for representative images, see Figure 1c). It features robust imaging, image analysis and data processing, as concluded from two parallel procedures carried out at independent institutions, the Department of Molecular Life Sciences at University of Zurich (UZH), and the Biomolecular Screening Facility at Ecole Polytechnique Fédérale de Lausanne (EPFL).

**Fig. 1:**
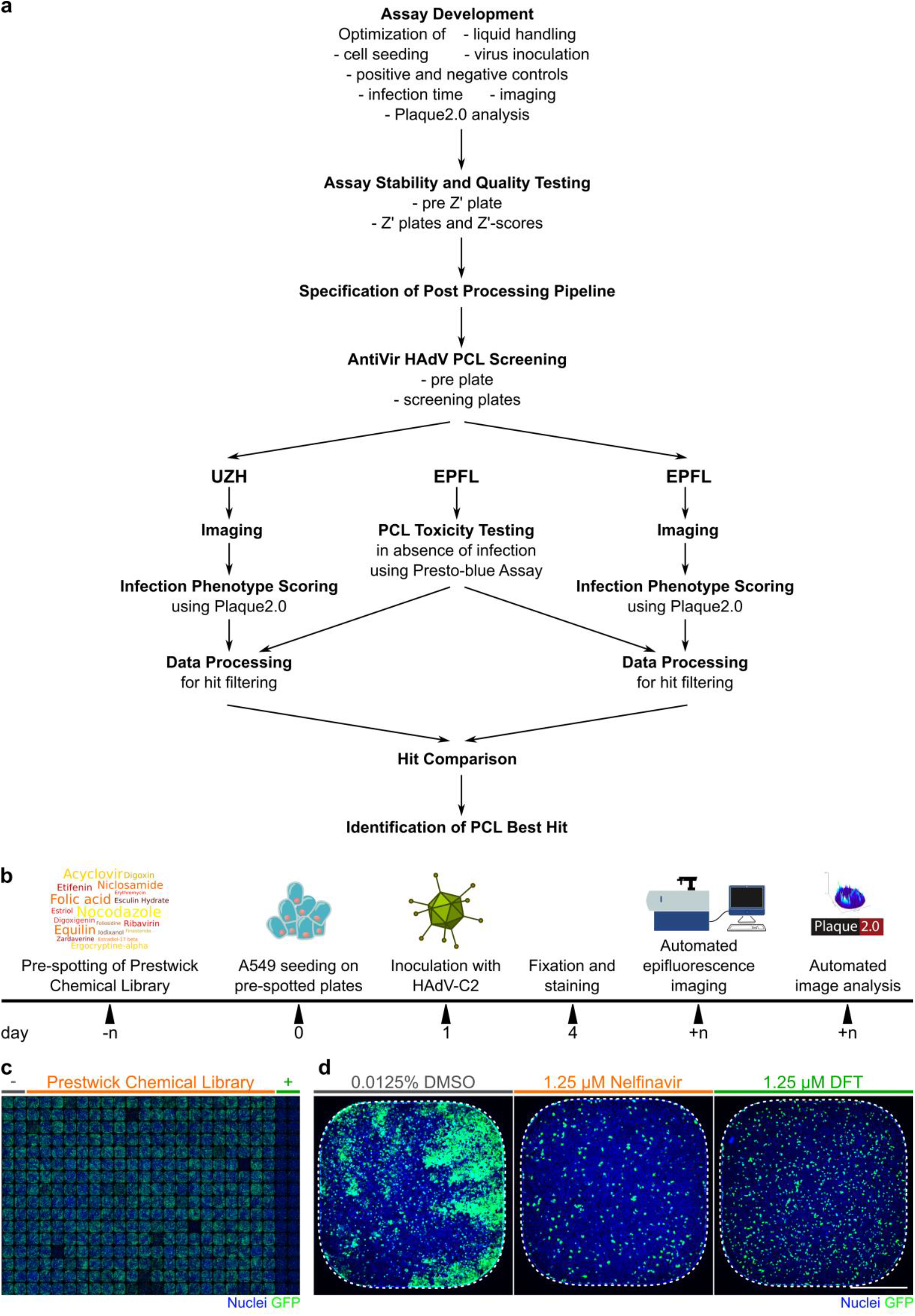
The compound screening procedure. **a** Following assay development, stability and quality testing, the screening of the PCL against HAdV infection was performed. Imaging, image analysis and data processing were independently carried out at UZH and EPFL, before hit ranking. **b** Schematic overview of the wet-lab pipeline. PCL compounds and DFT positive control in DMSO as well as DMSO alone as negative control were pre-spotted onto 384-well imaging plates by Echo acoustic liquid handling at 10 nl corresponding to a final concentration of 1.25 μM in 80 μl assay volume / well and stored at −20°C. Compound-blinded plates are thawed and 4,000 A549 cells / wells seeded. The following day, the cells were inoculated with HAdV-C2-dE3B at 1.77*10^5^ genome equivalents / well. Allowing for multiple viral replication rounds, the cells were PFA-fixed at 72 hpi and the nuclei stained with Hoechst 33342. The infection phenotypes were imaged using an epifluorescence HT microscope and scored using Plaque2.0. The data of the four technical replicates were further processed in R or through EPFL-BSF LIMS. **c** Exemplary epifluorescence microscopy images of cells in 384-wells stitched to a screening plate overview of 32 replicates of negative (two most left columns) and positive control (two most right columns) and 320 blinded PCL compounds (centre 20 columns). Hoechst-stained nuclei are shown in blue, viral GFP in green. **d** Representative 384-well epifluorescence microscopy images of the DMSO negative control (most left), the DFT positive control (most right) and the top hit Nelfinavir mesylate (centre). Hoechst-stained nuclei are shown in blue, virally expressed GFP in green. Scale bar is 5 mm.

Five phenotypic features were used to score the effects of the compounds on HAdV-C2-dE3B-GFP infected human lung cancer epithelial A549 cells – the number of infected and uninfected cell nuclei, the infection index (infected nuclei per total nuclei), the number of plaques (areas of infection foci originating from a single infected cell) and the integrated signal of the infection marker green fluorescence protein (GFP) encoded in the reporter virus genome. All data are available at the Image Data repository (IDR) ^1^, IDR accession number idr0081, and can be accessed via the IDR web client. Raw and scored infection phenotypes are provided for UZH and EPFL analyses. Rigorous assay development ensured a high assay quality, as indicated by mean Z’-factors of 0.52 for the plaque numbers. The screening was performed in four biological replicates at high reproducibility, and compounds that gave significant toxicity in uninfected cells were excluded during hit filtering. Imaging, image analysis and scoring by the two independent teams yielded well correlated scores and a congruent list of top hits, provided in Table 1. We confirmed the antiviral efficacy of Nelfinavir in a follow-up study (Georgi et al., in preparation).

**Tab. 1:**
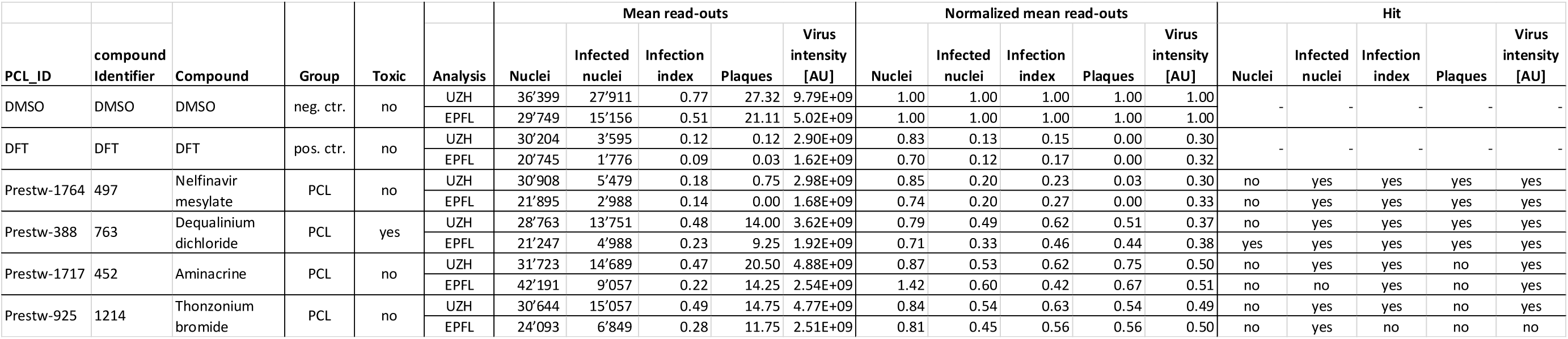
Summary of screening controls and top hits. Mean corresponds to the means over four biological replicates of PCL compound and 16 biological replicates each carrying 32 technical replicates for each control. Neg. ctr. refers to solvent control (DMSO), pos. ctr. to DFT-treated wells. Normalized indicates the mean read-outs of each compound relative to the mean of the positive control over all replicates. Toxicity was accessed by presto-blue assay of 3.5-day treatment of uninfected A549 cells as well as by the nuclei Z’-factor in the screen. Hits were selected for low toxicity and high inhibitory effects compared to solvent control samples. Note that compounds were scored toxic, if they showed significant toxicity in either of the assays.

## Methods

### Virus

HAdV-C2-dE3B-GFP was produced as described ^25^ and fully sequenced (GenBank accession number MT277585). In brief, the virus was generated by exchange of the viral E3B genome region with a reporter cassette harbouring the enhanced green fluorescent protein (GFP) under the immediate early Cytomegalovirus (CMV) promoter ^25^. The virus was grown in A549 cells and purified by double caesium chloride gradient centrifugation ^34^. Aliquots supplemented with 10% glycerol (v/v) were stored at −80°C. HAdV-C2-dE3B-GFP was found to be homogeneous by SDS-PAGE and negative-stain analyses by transmission electron microscopy.

### Cell culture

A549 (human adenocarcinomic alveolar basal epithelium) cells were obtained from the American Type Culture Collection (ATCC), Manassas, USA. The cells were maintained in full medium: high glucose Dulbecco Modified Eagle Medium (DMEM; Thermo Fisher Scientific, Waltham, USA) containing 7.5% fetal bovine serum (FBS, Invitrogen, Carlsbad, USA), 1% L-glutamine (Sigma-Aldrich, St. Louis, USA) and 1% penicillin streptomycin (Sigma-Aldrich, St. Louis, USA) and subcultured following PBS washing and trypsinisation (Trypsin-EDTA, Sigma-Aldrich, St. Louis, USA) weekly. Cells were grown at standard conditions (37°C, 5% CO2, 95% humidity) and passage number kept below 20.

### Preparation of pre-plates

Ten μl 0.0125% DMSO in PBS was spotted on all 384 wells each of imaging-compatible 384-well plates (Matrix plates #4332, Thermo Fisher Scientific, Waltham, USA) using a Matrix WellMate dispenser and normal bore Matrix WellMate tubing cartridges (Thermo Fisher Scientific, Waltham, USA). Plates were sealed and stored at −20°C.

### Blinding

The PCL compound arrangement as dispensed by EPFL in four subset plates A - D comprising each screening set replicate 1 - 4 was blinded and replaced by UZH with internal identifier (Supplementary Tables 2 and 3, *compoundidentifier* 1 to 1280). The identity of the compounds was only disclosed after the screening process and hit filtering (Supplementary Tables 2 and 3 and Table 1, *PCL_ID* Prestw-1 to Prestw-1804 and *compoundName)*.

### Compounds

The PCL was obtained from Prestwick Chemical (Illkirch, France). 3’-deoxy-3’-fluorothymidine (DFT, CAS number 25526-93-6) was obtained from Toronto Research Chemical, North York, Canada. All compounds were dissolved in dimethyl sulfoxide (DMSO, Sigma-Aldrich, St. Louis, USA) at a final stock concentration of 10 mM and stored at −20°C.

### Presto-blue toxicity assay

Toxicity of the PCL chemical compounds on uninfected A549 cells was tested using the Presto Blue Cell Viability reagent (Thermo Fisher Scientific, Waltham, USA). Briefly, following 3.5-day continuous treatment of A549 cells with compounds at concentrations and cell densities as in the screening protocol, 10% final PrestoBlue was added to each well and incubated for 1 h at standard cell incubation conditions. Fluorescence intensity (bottom-read) was measured using a multi-well plate reader (Tecan Infinite F500, Tecan, Männedorf, Switzerland) with excitation at 560/10 nm, emission at 590/10 nm at a fixed gain. Doxorubicin hydrochloride (Prestw-438, Prestwick Chemical, Illkirch, France) was used as a positive control for cytotoxicity, at a final concentration of 10 μM, and the corresponding concentration of the drug solvent DMSO was used as a negative control. The full PCL library was tested on duplicated plates. The EPFL-BSF in-house Laboratory Information Management System (LIMS) was used for data processing and statistical validation. First, raw PrestoBlue readings were normalized per plate to negative control values at 0 and positive controls at 1. Then, the normalized values of the duplicates were averaged. Assay quality was assessed for each plate through the Z’-factor calculation. Compounds were considered toxic when the normalized value for all replicates was higher than the average +3σ (standard deviation, SD) of the DMSO negative control for the corresponding plate. Scores and score SD were then calculated for hit compounds by averaging normalized value for all replicates.

### Preparation of plates for Z’-factor and drug screening

Ten nl of 10 mM PCL compounds, the nucleoside analogue DFT positive control (all dissolved in DMSO) and DMSO only as negative control were pre-spotted on imaging-compatible 384-well plates (Falcon plates, Corning Inc., New York, USA) using an Echo acoustic liquid handling system (Labcyte, San Jose, USA) by the EPFL-BSF, sealed and stored at −20°C. Each Z’-factor 384-well plate consisted of 192 technical replicates of positive and negative controls, each. Each screening plate set consisted of four subset plates A to D. Each screening plate comprised 32 technical replicates of positive and negative controls, each, and 320 PCL compounds.

### Wet-lab screening pipeline

The screening was performed in four independent biological replicates 1 - 4. Liquid handling was performed using a Matrix WellMate dispenser and Matrix WellMate tubing cartridges (Thermo Fisher Scientific, Waltham, USA). Prior to usage, tubings were rinsed with 125 ml autoclaved double-distilled (dd) H2O followed by 125 ml autoclaved PBS. Pre-spotted compound plates were thawed at room temperature (RT) for 30 min, briefly centrifuged before compounds were dissolved in 10 μl / well of PBS. 4,000 A549 cells / well in 60 μl full medium were seeded onto the compounds using standard bore tubing cartridges. Following cell adhesion over night, the cells were inoculated with 1.77*10^5^ genome equivalents per well of HAdV-C2-dE3B-GFP in 10 μl of full medium using bovine serum albumin (BSA, cell-culture grade, Sigma-Aldrich, St. Louis, USA)-blocked small bore tubing cartridges. The final compound concentration was 1.25 μM at 0.0125%

DMSO. Infection was allowed to progress over multiple infection rounds for 72 h giving rise to foc of infected cells originating from a single first round infected cell. Cells were fixed for 1 h at RT by addition of 26.6 μL 16% PFA and 4 μg/ml Hoechst 33342 (Sigma-Aldrich, St. Louis, USA) in PBS using standard bore tubing cartridges. Cells were washed three times with PBS before PBS supplemented with 0.02% N3 was added and plates were sealed for long-term storage at 4°C Following usage, tubings were rinsed with 125 ml autoclaved ddH2O followed by 125 m autoclaved PBS and autoclaved for re-usage.

### Imaging

Nuclei stained with Hoechst 33342 (DAPI channel) and viral GFP (FITC channel) were image on two devices. At UZH, plates were imaged on an IXM-C automated high-throughput fluorescence microscope (Molecular Devices, San Jose, USA) using MetaXpress (version 6.2 Molecular Devices, San Jose, USA) and a 4x air objective (Nikon S Fluor, 0.20 NA, 15.5 mm WD, Nikon Instruments, Minato, Japan) at widefield mode. Images of 2,048^2^ px at 1.72 μm/px resolution were acquired on an Andor sCMOS camera (Oxford Instruments, Abingdon, UK). Exposure times: DAPI 150 ms, FITC 20 ms. At EPFL, images were acquired on a IN Cell 2200 automated high-throughput fluorescence microscope (GE Healthcare, Chicago, USA) using IN Cell Analyzer (version 6.2, GE Healthcare, Chicago, USA) and a 4x air objective (Nikon Plan Apo 0.20 NA, 15.7 mm WD, Nikon Instruments, Minato, Japan) at widefield mode. Image size 2,048^2^ px at 1.625 μm/px resolution acquired on an Andor sCMOS camera. Exposure times: DAPI 300 ms, FITC 40 ms.

### Image analysis

The infection phenotype for each well was quantified by Plaque2.0 ^33^ (https://github.com/plaque2/matlab/tree/antivir) via five main read-outs: number of nuclei, number of infected nuclei, the ratio between infected and total nuclei referred to as infection index, number of multi-round infection foci termed plaques (plaque forming unit(s), pfu) and the integrated viral transgenic GFP intensity. Plaque2.0 parameters were optimized independently at UZH and EPFL for the data acquired at the respective institution.

### Z’-factor calculation

The Z’-factor was computed using R version 3.3.2 ^35^ according to Equation (1)

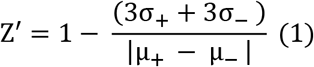

where σ_+_ is the SD of the positive control, σ_−_ is the SD of the negative control, μ_+_ the mean of the positive control and μ_−_ the mean of the negative control.

### Screening data processing

Plaque2.0 results were further processed and filtered. At UZH, results were processed in R version 3.3.2 ^35^, EPFL used KNIME version 3.4.0 ^36^ as well as the EPFL-BSF in-house LIMS. Mean infection scores over the Plaque2.0 read-outs of the four biological replicates of each PCL compound and the 16 biological replicates containing each 32 technical replicates of positive and negative control, each, were calculated. Each compound’s scores were normalized by the mean score of the DMSO negative control of the respective plate. Only non-toxic, effective PCL compounds were considered as HAdV inhibitor candidates. Non-toxic compounds were filtered by applying an inclusive μ_+_ (mean of the negative control) ± 2σ (SD of the negative control) threshold for number of nuclei. Efficacy was filtered by applying an excluding μ+ ± 3σ threshold for the infection scores (number of infected nuclei, infection index, number of plaques or integrated GFP intensity). Subsequently, compounds exhibiting significant toxicity to noninfected cells were excluded.

### Data Records

#### Data structure and repository

The screening data comprise the information collected during assay development, including stability, quality and screening of the PCL itself. The latter two were imaged on two different microscopes. We provide the parameters used for Plaque2.0 image analysis, and the code for the subsequent hit filtering in R. The data structure as available for download at the IDR ^1^, accession number idr0081, outlined in Figure 2a. Moreover, the data can be viewed conveniently on the IDR web client, where it is structured as outlined in Figure 2b.

**Fig. 2.**
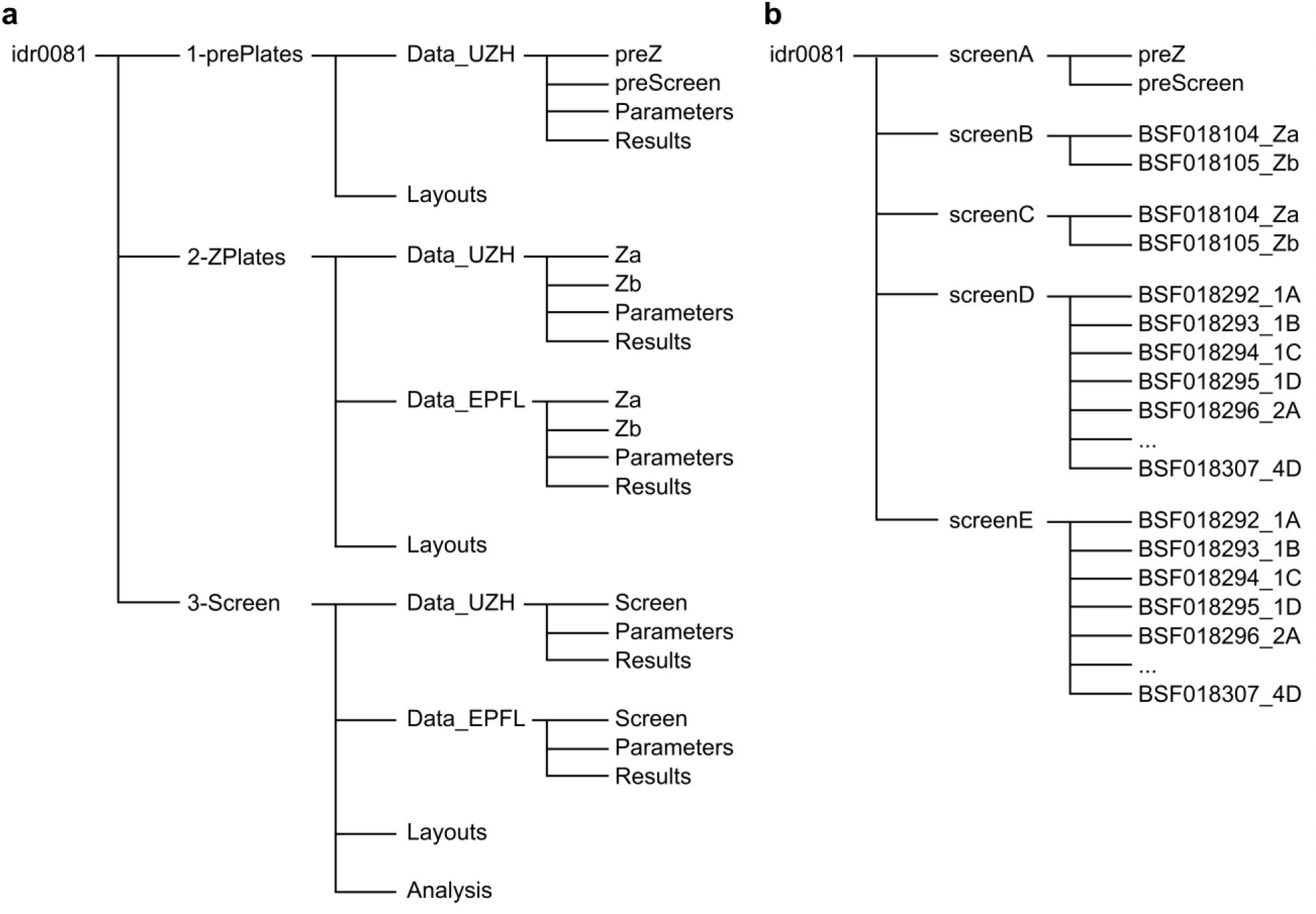
Project data structure available at IDR, accession number idr0081. **a** In the data provided for download, there are three sub-folders for *1-prePlates*, *2-ZPlates* and *3-Screen*. The latter two contain both the images and analyses generated by UZH and EPFL. **b** The data provided for viewing are divided into five screens: *screenA* contains the pre-plates and *screenB* and *screenC* consist of the Z’-factor plates imaged and analysed at UZH and EPFL, respectively. *screenD* and *screenE* provide the screening data obtained at UZH and EPFL, respectively.

#### Data sets and file types

The data provided for download consists of three data sets 1 to 3 (see Figure 2a).

*1-prePlates* contains layouts (.csv), images (.tif), Plaque2.0 image analysis parameters (.mat) and results (.csv) for the assay stability test plates performed at UZH prior to Z’-factor plates (*preZ*) and the screen (*preScreen*).
*2-ZPlates* contains layouts (.csv), images (.tif), Plaque2.0 image analysis parameters (.mat) and results (.csv) for the two Z’-factor plates *a* and *b* as imaged and analysed at UZH (*Data_UZH*) and EPFL (*Data_EPFL*).
*3-Screen* contains layouts (.csv), images (.tif), Plaque2.0 image analysis parameters (.mat) and results (.csv) for the 16 screening plates (four biological replicas *1 - 4*, each consisting of a set of four subset plates *A - D*) as imaged and analysed at UZH (*Data_UZH*) and EPFL (*Data_EPFL*). Moreover, *Analysis* contains the Plaque2.0 batch processing (*AntiVir_batchprocessing.m*) and hit filtering pipeline (*AntiVir_hitfiltering.R*) used by UZH. *Analysis* also contains the Presto-blue raw results (.csv) for toxicity in absence of infection.

The data provided for browsing via the IDR web client consists of five screens *A* to *E* (see Figure 2b).

idr0081-study.txt summarizes the overall study and the five screens that were performed.
*screenA* contains the assay stability test plates performed at UZH prior to Z’-factor plates (*preZ*) and the screen (*preScreen*). *idr0081-screenA-library.txt* provides thorough information on the tested compounds including PubChem identifiers and their plate layout. *idr0081-screenA-processed.txt* presents the results of the Plaque2.0-based image analysis. *idr0081-screenA-mean.txt* summarises the infection scores per pre plate.
*screenB* contains the assay quality test plates (Z’-factor plates *a* and *b*) performed at UZH. *idr0081-screenB-library.txt* provides thorough information on the tested compounds including PubChem identifiers and their plate layout. *idr0081-screenB-processed.txt* presents the results of the Plaque2.0-based image analysis. *idr0081-screenB-mean.txt* summarises the infection scores per Z’-factor plate.
*screenC* contains the assay quality test plates (Z’-factor plates *a* and *b*) performed at EPFL. *idr0081-screenC-library.txt* provides thorough information on the tested compounds including PubChem identifiers and their plate layout. *idr0081-screenC-processed.txt* presents the results of the Plaque2.0-based image analysis. *idr0081-screenC-mean.txt* summarises the infection scores per Z’-factor plate.
*screenD* contains the PCL screening plates (in replicates *1* to *4*, consisting of subset plates *A* to *D*) performed at UZH. *idr0081-screenD-library.txt* provides thorough information on the tested compounds including PubChem identifiers and their plate layout. *idr0081-screenD-processed.txt* presents the results of the Plaque2.0-based image analysis. *idr0081-screenB-filtered.txt* summarises the infection scores per compound and indicates if it was identified as hit.
*screenE* contains the PCL screening plates (in replicates *1* to *4*, consisting of subsets *A* to *D*) performed at EPFL. *idr0081-screenE-library.txt* provides thorough information on the tested compounds including PubChem identifiers and their plate layout. *idr0081-screenE-processed.txt* presents the results of the Plaque2.0-based image analysis. *idr0081-screenE-filtered.txt* summarises the infection scores per compound and indicates if it was identified as hit.

### Technical Validation

#### Assay stability

The wet-lab screening pipeline was optimized regarding liquid handling, cell seeding, virus inoculum, positive and negative controls, infection time, as well as imaging and image analysis. This ensured a high assay stability and reproducibility. Furthermore, all compounds, especially media and supplements, the BSA for tubing saturation, PFA- and Hoechst-supplemented fixative were prepared as large batch from a single lot and stored as single-use aliquots. Prior to every experiment, assay stability with respect to cell and infection phenotype was tested on pre-plates according to the established wet-lab, imaging and image analysis pipeline. Since the solvent control had already been spotted in 10 μl PBS, no further PBS was added prior to cell seeding. Periodically, the virus stock dilution was tested and adjusted for experiments if necessary.

#### Assay quality determination: Z’-factor

The accuracy of the wet-lab, imaging and image analysis pipeline was assessed by two independently imaged and analyzed Z’-factor plates (Table 2 and Figure 3). 3σ Z’-factors of *numberOfInfectedNuclei, infectionIndex* and *numberOfPlaques* were in the range of 0.30 to 0.57, scoring good to excellent. *totalVirusIntensity* (Z’-factors between −0.07 to 0.08) were not suitable to identify HAdV infection inhibitors, while *numberOfNuclei* (Z’-factors between −1.11 to −8.10) was not a useable readout either. Additionally, the Z’-factors were determined for each of the 16 screening plates (Table 3 and Figure 4). 3σ Z’-factors of *numberOfInfectedNuclei*, *infectionIndex* and *numberOfPlaques* were in the range of 0.27 to 0.57, scoring good to excellent.

**Fig. 3:**
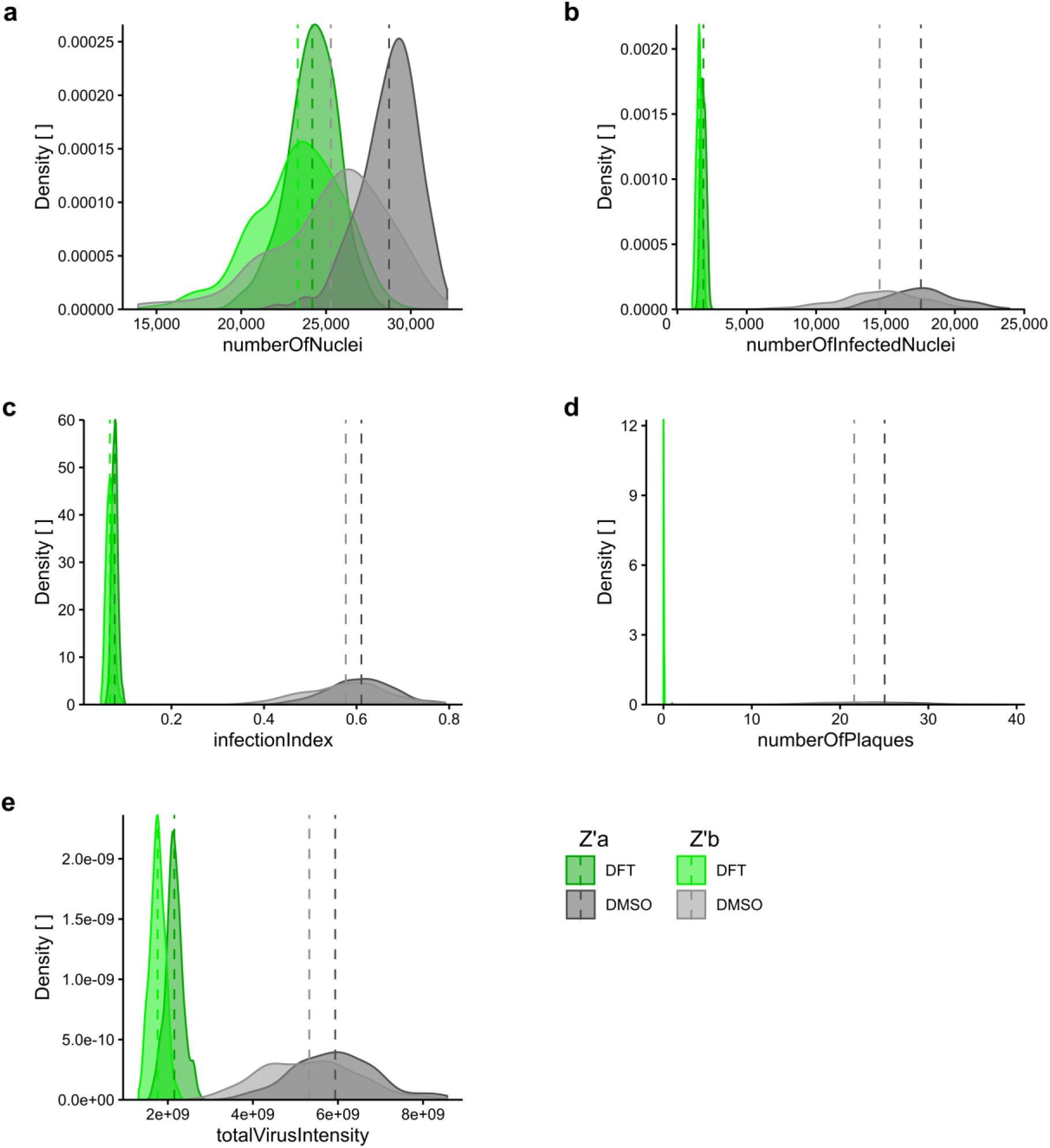
Infection score density of positive and negative controls across Z’-factor plates. Distribution of **a** *numberOfNuclei*, **b** *numberOfInfectedNuclei*, **c** *infectionIndex*, **d** *numberOfPlaques* and **e** *totalVirusIntensity* in negative control (0.0125% DMSO) compared to positive control-treated (1.25 μM DFT) samples of the two Z’-factor plates. Dark green and dark grey indicate Z’-factor plate a, light green and grey show Z’-factor plate b. Dashed vertical lines mark mean of 192 technical replicates.

**Fig. 4:**
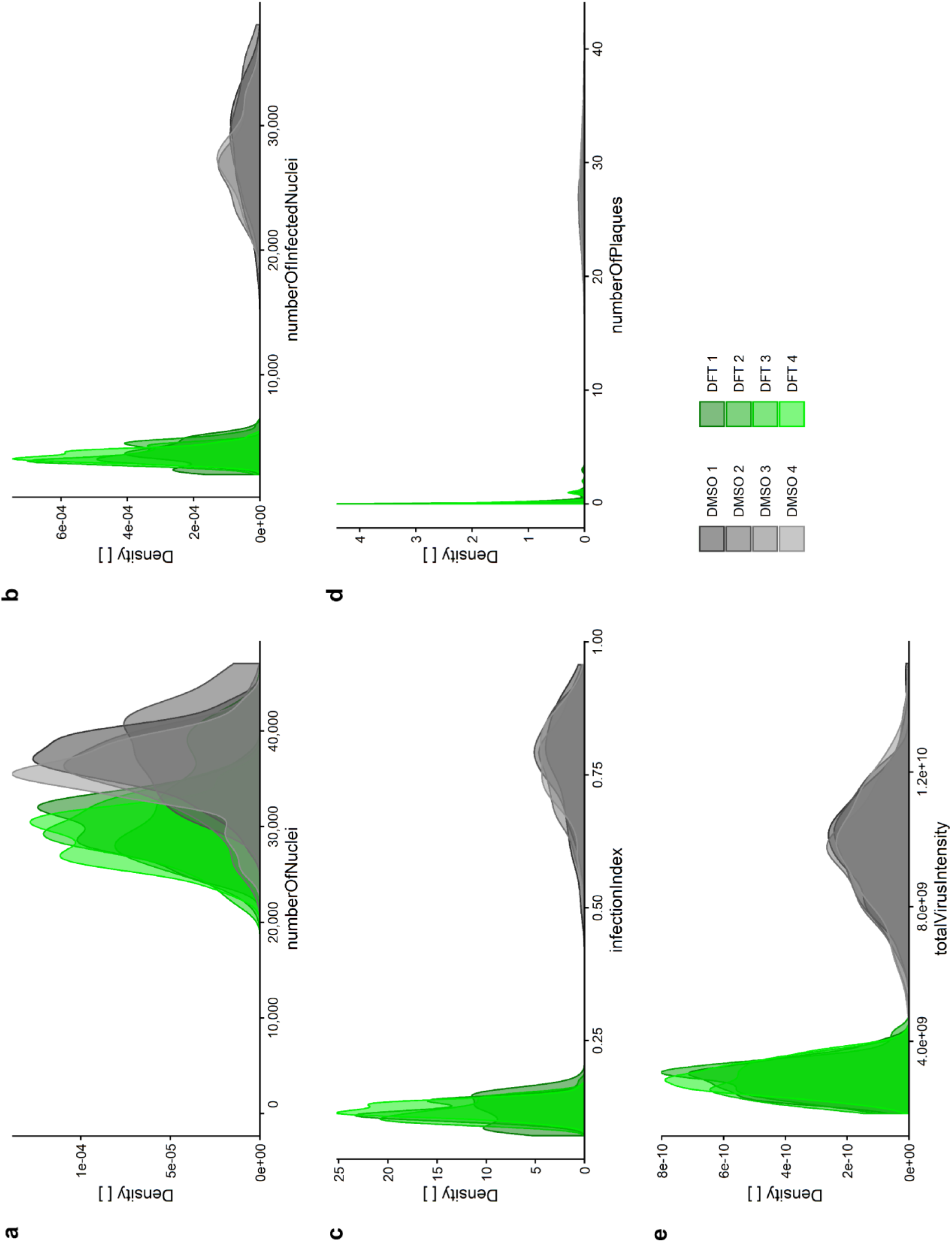
Infection score density of positive and negative controls across screening replicates. Distribution of **a** *numberOfNuclei*, **b** *numberOfInfectedNuclei*, **c** *infectionIndex*, **d** *numberOfPlaques* and **e** *totalVirusIntensity* in negative control (0.0125% DMSO in grey) compared to positive control-treated (1.25 μM DFT in green) samples of the screening sets. Each replicate 1 to 4 indicated by colour shading is comprised of four plates containing 32 technical replicas per control.

**Tab. 2:** Z’-factor plates. The quality of the screening platform was assessed prior to screening of the PCL by two independent Z’-factor plates containing 192 technical replicates of both positive control (1.25 μM DFT) and solvent only control (0.0125% DMSO). Z’-factors for the five Plaque2.0 read-outs ^33^ obtained by independent analysis at UZH and EPFL were calculated according to Equation (1) for 3 and 2σ.

| Barcode | Plate | UZH |  |  |  |  | EPFL |  |  |  |  |
| --- | --- | --- | --- | --- | --- | --- | --- | --- | --- | --- | --- |
|  |  | 3 sigma |  |  |  |  | 3 sigma |  |  |  |  |
|  |  | numberOf Nuclei | numberOf Infected Nuclei | infection Index | numberOf Plaques | totalVirus Intensity | numberOf Nuclei | numberOf Infected Nuclei | infection Index | numberOf Plaques | totalVirus Intensity |
| BSF018104 | Za | -1.11 | 0.50 | 0.57 | 0.50 | 0.07 | -1.20 | 0.36 | 0.47 | 0.52 | 0.08 |
| BSF018105 | Zb | -8.10 | 0.30 | 0.45 | 0.39 | -0.07 | -1.23 | 0.27 | 0.32 | 0.44 | -0.04 |
| Mean |  | -4.61 | 0.40 | 0.51 | 0.44 | 0.00 | -1.22 | 0.32 | 0.40 | 0.48 | 0.02 |
| Barcode | Plate | UZH |  |  |  |  | EPFL |  |  |  |  |
|  |  | 2 sigma |  |  |  |  | 2 sigma |  |  |  |  |
|  |  | numberOf Nuclei | numberOf Infected Nuclei | infection Index | numberOf Plaques | totalVirus Intensity | numberOf Nuclei | numberOf Infected Nuclei | infection Index | numberOf Plaques | totalVirus Intensity |
| BSF018104 | Za | -0.41 | 0.67 | 0.71 | 0.67 | 0.07 | -0.47 | 0.58 | 0.64 | 0.68 | 0.38 |
| BSF018105 | Zb | -5.07 | 0.53 | 0.63 | 0.59 | -0.07 | -0.49 | 0.52 | 0.55 | 0.63 | 0.31 |
| Mean |  | -2.74 | 0.60 | 0.67 | 0.63 | 0.00 | -0.48 | 0.55 | 0.60 | 0.66 | 0.35 |

**Tab. 3:** Z’-factors of screening plates. The quality of the screening data was assessed for each screening plate based on the 32 technical replicates of both positive control (1.25 μM DFT) and solvent only control (0.0125% DMSO) in each plate. Z’-factors for the five Plaque2.0 read-outs ^33^ obtained by independent analysis at UZH and EPFL were calculated according to Equation (1) for 3σ.

| Barcode | Plate | UZH |  |  |  |  | EPFL |  |  |  |  |
| --- | --- | --- | --- | --- | --- | --- | --- | --- | --- | --- | --- |
|  |  | 3 sigma |  |  |  |  | 3 sigma |  |  |  |  |
|  |  | numberOf<br>Nuclei | numberOf<br>Infected<br>Nuclei | infection<br>Index | numberOf<br>Plaques | totalVirus<br>Intensity | numberOf<br>Nuclei | numberOf<br>Infected<br>Nuclei | infection<br>Index | numberOf<br>Plaques | totalVirus<br>Intensity |
| BSF018292 | 1A | -0.13 | 0.58 | 0.58 | 0.59 | 0.35 | -0.14 | 0.51 | 0.49 | 0.58 | 0.31 |
| BSF018293 | 1B | -0.88 | 0.58 | 0.65 | 0.55 | 0.34 | -0.35 | 0.51 | 0.52 | 0.51 | 0.35 |
| BSF018294 | 1C | -1.01 | 0.62 | 0.62 | 0.63 | 0.33 | -0.74 | 0.52 | 0.50 | 0.66 | 0.32 |
| BSF018295 | 1D | -0.34 | 0.56 | 0.54 | 0.45 | 0.16 | -0.21 | 0.43 | 0.38 | 0.46 | 0.19 |
| BSF018296 | 2A | -1.35 | 0.64 | 0.67 | 0.55 | 0.30 | -0.20 | 0.57 | 0.55 | 0.55 | 0.28 |
| BSF018297 | 2B | -3.63 | 0.56 | 0.52 | 0.45 | 0.14 | -1.20 | 0.45 | 0.39 | 0.40 | 0.12 |
| BSF018298 | 2C | -1.81 | 0.60 | 0.58 | 0.49 | 0.24 | -0.38 | 0.52 | 0.42 | 0.52 | 0.19 |
| BSF018299 | 2D | -1.94 | 0.57 | 0.57 | 0.57 | 0.24 | -0.22 | 0.50 | 0.43 | 0.63 | 0.20 |
| BSF018300 | 3A | -1.74 | 0.64 | 0.66 | 0.56 | 0.36 | -0.54 | 0.55 | 0.51 | 0.59 | 0.34 |
| BSF018301 | 3B | -1.13 | 0.60 | 0.68 | 0.58 | 0.40 | -0.09 | 0.52 | 0.57 | 0.59 | 0.40 |
| BSF018302 | 3C | -4.02 | 0.66 | 0.68 | 0.48 | 0.42 | -1.07 | 0.63 | 0.60 | 0.50 | 0.41 |
| BSF018303 | 3D | -2.36 | 0.55 | 0.63 | 0.51 | 0.36 | -0.10 | 0.58 | 0.54 | 0.52 | 0.35 |
| BSF018304 | 4A | -0.68 | 0.70 | 0.74 | 0.42 | 0.37 | -0.29 | 0.56 | 0.58 | 0.48 | 0.36 |
| BSF018305 | 4B | -0.17 | 0.71 | 0.74 | 0.51 | 0.50 | -0.50 | 0.63 | 0.67 | 0.50 | 0.50 |
| BSF018306 | 4C | -0.44 | 0.61 | 0.62 | 0.50 | 0.28 | -0.28 | 0.50 | 0.48 | 0.47 | 0.26 |
| BSF018307 | 4D | -0.77 | 0.63 | 0.70 | 0.42 | 0.41 | -0.22 | 0.54 | 0.56 | 0.36 | 0.39 |
| Mean |  | -1.40 | 0.61 | 0.64 | 0.52 | 0.32 | -0.41 | 0.53 | 0.51 | 0.52 | 0.31 |

#### Independent analysis and filtering

Imaging, image analysis and screening data processing were performed by two independent research teams at UZH and EPFL, as depicted in Figure 1. Raw and scored infection phenotypes are shown for UZH and EPFL analyses (Supplementary Tables 2, 3 and Supplementary Tables 4, 5, respectively). Both dry-lab pipelines confirmed the high assay quality (Tables 2 and 3). During hit filtering, PCL compounds that gave significant toxicity in uninfected cells were excluded during hit filtering (Figure 5, Table 4). As summarized in Figure 6 left panel, both scores are strongly correlated with R^2^ between 0.6870 - 0.9870. Both approaches yielded identical top scored compounds (Figure 6, right panel), of which Prestw-1764, Nelfinavir mesylate, was the top hit.

**Fig. 5:**
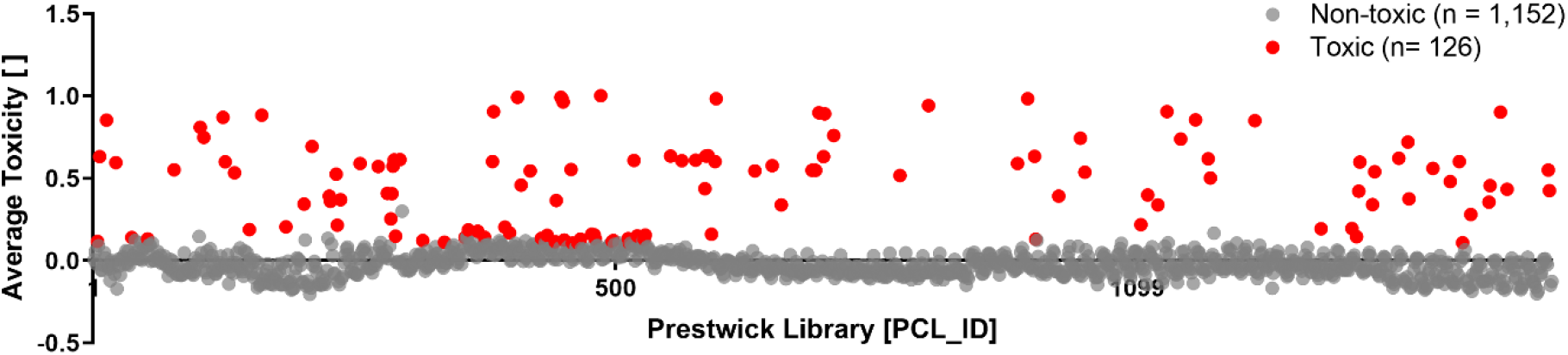
PCL compound toxicity in uninfected cells. Of the 1,278 PCL compounds tested, 126 PCL compounds are found to be toxic, as shown in red, and listed in Table 4. A549 cells were treated with PCL compounds in duplicates according to the screening wet-lab protocol, however, in absence of HAdV infection for 3.5 days. Doxorubicin hydrochloride (Prestw-438) was used as a positive control for cytotoxicity, at a final concentration of 10 μM, and the corresponding concentration of the drug solvent DMSO was used as a negative control. Cell viability was determined by Presto-blue assay. Presto-blue fluorescence intensities of each well were normalized per plate to negative control values at 0 and positive controls at 1. Compounds were considered toxic, when the normalized value for all replicates was higher than the average +3σ (standard deviation, SD) of the DMSO negative control for the corresponding plate. X-axis indicates compounds by their PCL identifier (*PCL ID*, see Supplementary Table 1). Normalized average presto-blue read-outs are depicted on the y-axis.

**Tab. 4:**
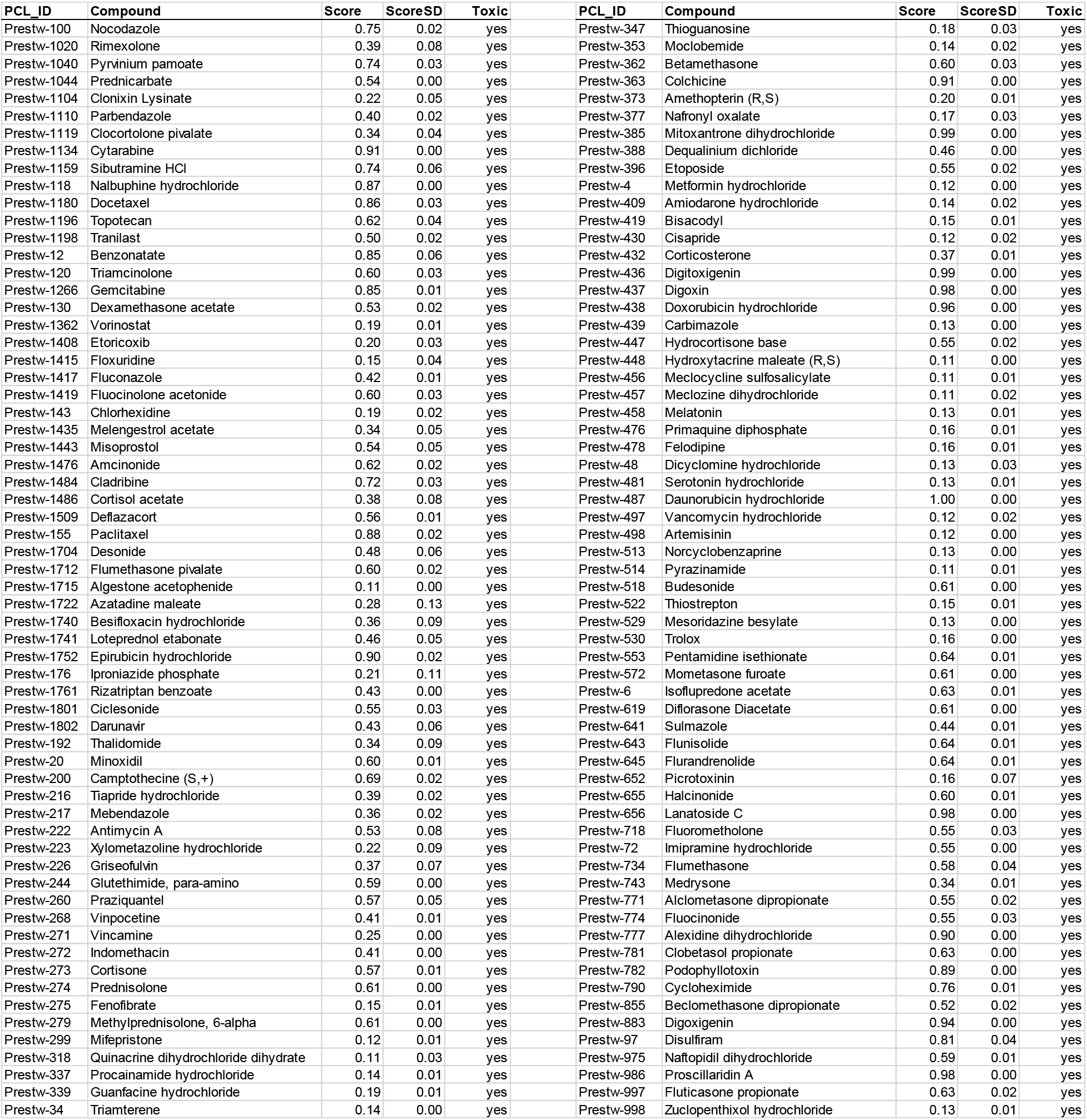
PCL compounds excluded due to toxicity in uninfected cells. Presto-blue raw data are available at *idr0081/3-Screen/Analysis/Toxicity.xls*.

**Fig. 6:**
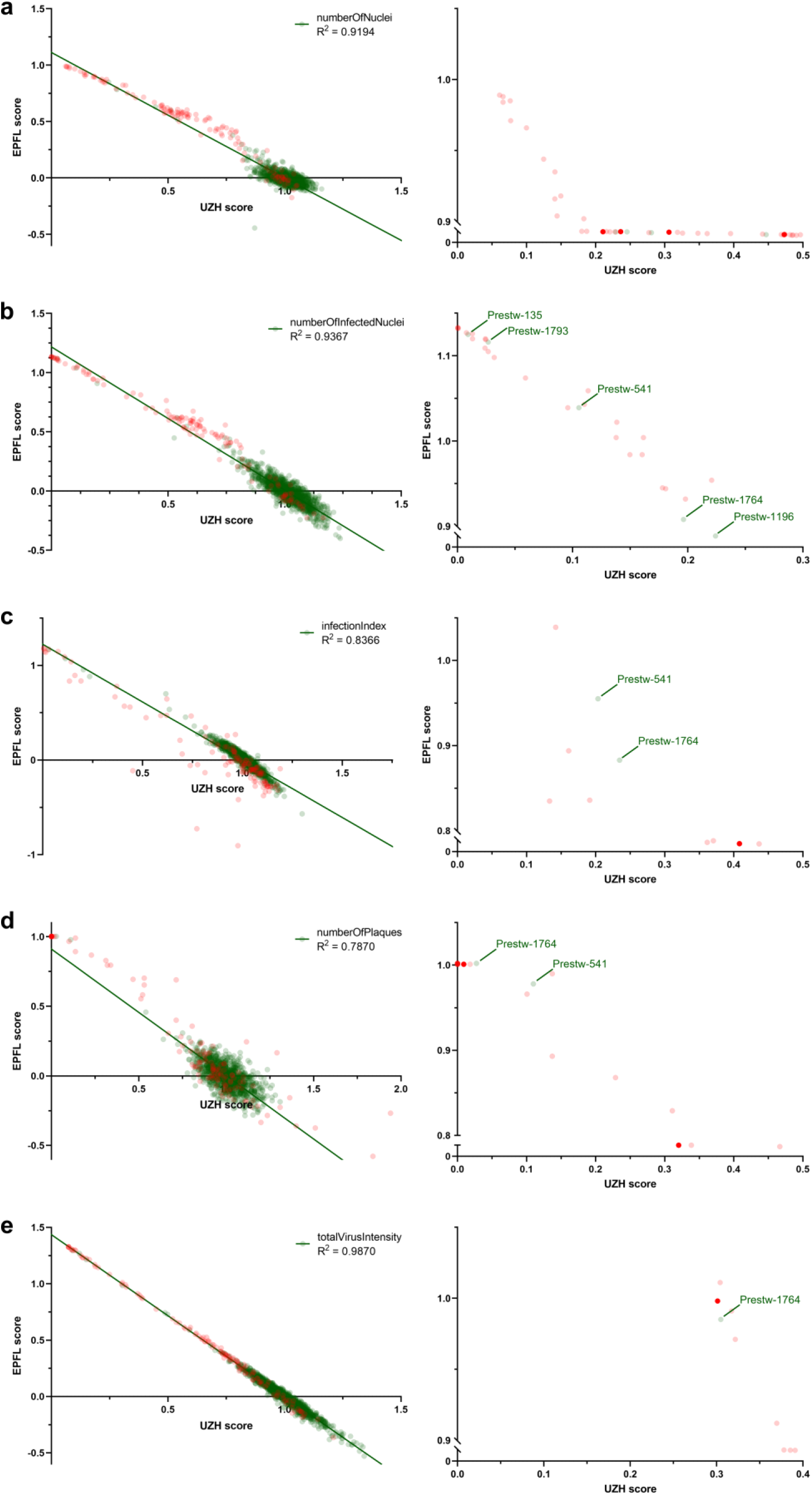
Correlation between scores from independent dry-lab pipelines. Imaging, image analysis and data processing is performed independently at UZH and EPFL. PCL-treated infection phenotypes from 4 biological replicates were averaged and normalized against the DMSO solvent control. Obtained scores for **a** *numberOfNuclei*, **b** *numberOfInfectedNuclei*, **c** *infectionIndex*, **d** *numberOfPlaques* and **e** *totalVirusIntensity* of the 1,278 tested PCL compounds from UZH and EPFL are correlated via linear regression (green line), R^2^ is calculated using GraphPad Prism 8.2.1. Highest scoring compounds are shown on the right and *PCL_ID* of non-toxic compounds indicated. Red dots indicate toxicity in the absence of infection, non-toxic compounds are shown in green.

### Usage Notes

Five parameters were used to score the infection phenotype of each well: the number of nuclei (*numberOfNuclei*), number of infected nuclei (*numberOfInfectedNuclei*), the ratio between number of infected and total nuclei (*infectionIndex*), the number of multi-round infection foci termed plaques (*numberOfPlaques*) and the extend of viral GFP reporter expression as integrated GFP intensity *totalVirusIntensity*).

#### Infection scoring using the Plaque2.0 GUI

A detailed manual for Plaque2.0 GUI-based infection phenotype scoring is available at *plaque2.github.io/*. No MATLAB license is necessary.

The following settings should be used:

##### Input/Output

*Processing Folder*. Path to folder containing the images (e.g. *idr0081/3-Screen/Data_EPFL/Screen/ BSF018292_ 1A*).

*filename pattern* Data_UZH: *.* (?<wellName>[A-Z][0-9]*)_(?<channelName>w[0-9]*).TIF*

*filename pattern* Data_EPFL: *.* (?<wellName>[A-Z] - [0-9]+)[(]fld 1 wv (?<channel>[A-Z]{4}) *.tif*

*Plate name:* Name of the plate to be analysed (e.g. *BSF018292_1A*)

*Result Output Folder*. Path to the results folder in the respective data folder (e.g. *idr0081/3-Screen/Data_EPFL/Results*).

##### Stitch

Stitching of the images is not necessary, since every 384-well is imaged in a single site. Do not activate the tab.

##### Mask

*Custom Mask File:* Path to the manually defined mask file (e.g. *idr0081/3-Screen/Data_UZH/Parameters*). Masking is optional and was not performed by EPFL.

##### Monolayer

*Channel:* Nuclei were imaged in channel 1.

##### Plaque

*Channel*: Viral GFP reporter signal was imaged in channel 2.

#### Infection scoring using the Plaque2.0 batch script

How to use the *AntiVir_batchprocessing.m* for Plaque2.0 batch processing is indicated in the comments of the code.

### Code Availability

#### Plaque2.0 batch image analysis for infection scoring

The MATLAB (version R2016b, The MathWorks, Natick, USA) script *AntiVir_batchprocessing.m* used by UZH for image analysis is provided for download at IDR, accession number idr0081, under *idr0081/3-Screen/Analysis*. It is based on the Plaque2.0 software available on GitHub under GPLv3 open source license: https://github.com/plaque2/matlab.

To batch analyse the HAdV screening data by Plaque2.0, fork or download the Plaque2.0 AntiVir code from GitHub: https://github.com/plaque2/matlab/tree/antivir. Place the *AntiVir_batchprocessing.m* file from *idr0081/3-Screen/Analysis* into the *Plaque2/matlab* folder and follow the instructions in *AntiVir_batchprocessing.m*. A MATLAB license is required.

#### Hit filtering using R

The R^35^ (version 3.6.1 (2019-07-05)) script *AntiVir_hitfiltering.R* used by UZH for data processing and hit filtering is provided at IDR accession number idr0081 under *idr0081/3-Screen/Analysis*.

## Author Contributions

UFG, VA, AY conceived the screening idea. FG designed the experiments, and with UFG coordinated the project. FK prepared the PCL-spotted plates. FG and RW performed the experiments. FG and FK acquired the data. FG and VA analysed the imaging data. LM and FG processed the data. GT organized and supervised the screening project at the EPFL-BSF. FG, FK and UFG wrote manuscript, with input from all the co-authors.

## Acknowledgements

We thank the entire Greber lab for fruitful discussions and critical assessment of the data. We further thank the IDR team for making our work openly accessible.

## Competing Interests

The authors declare no conflict of interest.

## Funding

The work was supported by the Swiss National Science Foundation to UFG (Grant numbers 316030_170799 / 1 and 31003A_179256 / 1), and the SNSF through the National Research Program “NCCR chemical biology” to GT and UFG.

## Abbreviations

BSA: bovine serum albumin
BSF: Biomolecular Screening Facility
CMV: Cytomegalovirus
DFT: 3’-Deoxy-3’-fluorothymidine
DMEM: Dulbecco’s Modified Eagle medium
DMSO: Dimethyl sulfoxide
dpi: days post infection
EPFL: Ecole Polytechnique Fédérale de Lausanne
FBS: fetal bovine serum
GFP: green fluorescent protein
HAdV: Human adenovirus
hpi: hours post infection
HTS: high-throughput screening
IDR: The Image Data Resource
LIMS: Laboratory Information Management System
LUT: Look up table
PCL: Prestwick Chemical Library
PFA: para-formaldehyde
pfu: plaque forming unit(s)
RT: room temperature
SE: standard error
SD: standard deviation
UZH: University of Zurich

---

**Supplementary Table 1: PCL compounds tested in the screening procedure.**

PCL catalogue IDs (*PCL_ID*), compound names (*CompoundName*), PubChem identifier (*CompoundPubChemCID*) and link (*CompoundPubChemURL*), the tested concentration in μM (*CompoundConcentrationMicroMolar*), the CAS registry number (*CAS*), structure according SMILES notation (*CompoundSMILES*), acoustic dispensing spottability (*SpottabilityFlag*) and group (*Group*) for each of the 1,280 PCL compounds and control compounds. Two compounds, Prestw-354 (Clopamide) and Prestw-410 (Amphotericine B) could not be successfully transferred via acoustic dispensing due to precipitation, and were not included in the screening.

**Supplementary Table 2: Raw Plaque-2.0 infection scores of the HAdV PCL screening imaged and analysed at UZH.**

*virus* indicates virus genotype, the PCL was tested against, *compoundIdentifier* indicates the UZH identifier for blinded testing by UZH, *setPlate* is the subset plate A to D and *replicate* refers to the replicate 1 to 4, *wellRow* and *wellColumn* indicate the well and *plate* indicate the screening plate sequence number. The Plaque2.0-based infection scores are *numberOfNuclei* reporting the number of nuclei based on Hoechst staining, *numberOfInfectedNuclei* refers to the number of GFP reporter-based number of infected nuclei, *infectionIndex* is the ratio of *numberOfInfectedNuclei* to *numberOfNuclei*, the number of GFP reporter-based plaques is given by *numberOfPlaques* and *totalVirusIntensity* indicates total GFP reporter signal intensity.

**Supplementary Table 3: Processed Plaque-2.0 infection scores of the HAdV PCL screening imaged and analysed at UZH.**

*virus* indicates virus genotype, the PCL was tested against, *compoundIdentifier* indicates the UZH identifier for blinded testing by UZH, *PCL_ID* and *compoundName* disclose the PCL compound identifier and name, respectively. *Barcode1, Barcode2, Barcode3* and *Barcode4* indicate on which screening plates, given by the screening plate sequence number defined by EPFL, the PCL compound was tested on. The Prestoblue toxicity scoring of the compound tested in noninfected cells is given as *1* (toxic) and *0* (non-toxic) in *nonInfectedToxHit*. The mean Plaque2.0-based infection scores of the four biological replicates are provided by *mean_numberOfNuclei* (number of nuclei based on Hoechst staining), *mean_numberOfInfectedNuclei* (number of GFP reporter-based number of infected nuclei), *mean_infectionIndex* (ratio of *numberOfInfectedNuclei* to *numberOfNuclei*), *mean_numberOfPlaques* (number of GFP reporter-based plaques) and *mean_totalVirusIntensity* (total GFP reporter signal intensity). The infection scores of the positive and negative controls are averaged (mean) over the 32 technical replicates, each, per plate, and the mean PCL compound infection scores were normalized by the mean negative control infection score of the respective plate indicated by by *mean_numberOfNucleiRel* (number of nuclei based on Hoechst staining), *mean_numberOfInfectedNucleiRel* (number of GFP reporter-based number of infected nuclei), *mean_infectionIndexRel* (ratio of *numberOfInfectedNuclei* to *numberOfNuclei), mean_numberOfPlaquesRel* (number of GFP reporter-based plaques) and *mean_totalVirusIntensityRel* (total GFP reporter signal intensity).

**Supplementary Table 4: Raw Plaque-2.0 infection scores of the HAdV PCL screening imaged and analysed at EPFL.**

*Barcode* indicates the screening plate sequence number defined by EPFL and *Well Position* gives the well. Plaque2.0-based infection scores are *numberOfNuclei* reporting the number of nuclei based on Hoechst staining, *numberOfInfectedNuclei* refers to the number of GFP reporter-based number of infected nuclei, *infectionIndex* is the ratio of *numberOfInfectedNuclei* to *numberOfNuclei*, the GFP reporter-based number of plaques is given by *numberOfPlaques* and *totalVirusIntensity* indicates total GFP reporter signal intensity.

**Supplementary Table 5: Processed Plaque-2.0 infection scores of the HAdV PCL screening imaged and analysed at EPFL.**

*Name* indicates the name of the tested PCL compound. The Plaque2.0-based infection scores of the four biological replicates of each PCL compound were averaged (mean). The Plaque2.0-based infection scores of the positive and negative controls are averaged (mean) over the 32 technical replicates, each, per plate. Each compound’s scores were normalized by the mean score of the negative control of the respective plate and indicated by *Mean N_nuclei* (number of nuclei based on Hoechst staining), *Mean N_infected* (number of GFP reporter-based number of infected nuclei), *Mean InfIndex* (ratio of *numberOfInfectedNuclei* to *numberOfNuclei*), *Mean N_plaques* (number of GFP reporter-based plaques) and *Mean TotVirInt* (total GFP reporter signal intensity). Non-toxic compounds were filtered by applying an inclusive μ+ (mean of the negative control) ± 2σ (SD of the negative control) threshold for number of nuclei. Efficacy was filtered by applying an excluding μ+ ± 3σ (SD of the negative control) threshold for the infection scores. The obtained scores for each infection score of each PCL compound indicated as *Mean Scores N_nuclei* (number of nuclei based on Hoechst staining), *Scores N_Infected* (number of GFP reporter-based number of infected nuclei), *Scores InfIndex* (ratio of *numberOfInfectedNuclei* to *numberOfNuclei), Scores N_plaques* (number of GFP reporter-based plaques) and *Scores TotVirInt* (total GFP reporter signal intensity). Subsequently, compounds exhibiting significant toxicity to noninfected cells were excluded.

## Notes

https://idr.openmicroscopy.org/webclient/

### Competing Interest Statement

The authors have declared no competing interest.

### Summary of Updates

We received the GenBank accession number MT277585 and indicate the data structure for the IDR web client.

