## Supplementary figures and images for "High-content image-based drug screen identifies a clinical compound against cell transmission of adenovirus"

### Figure 1

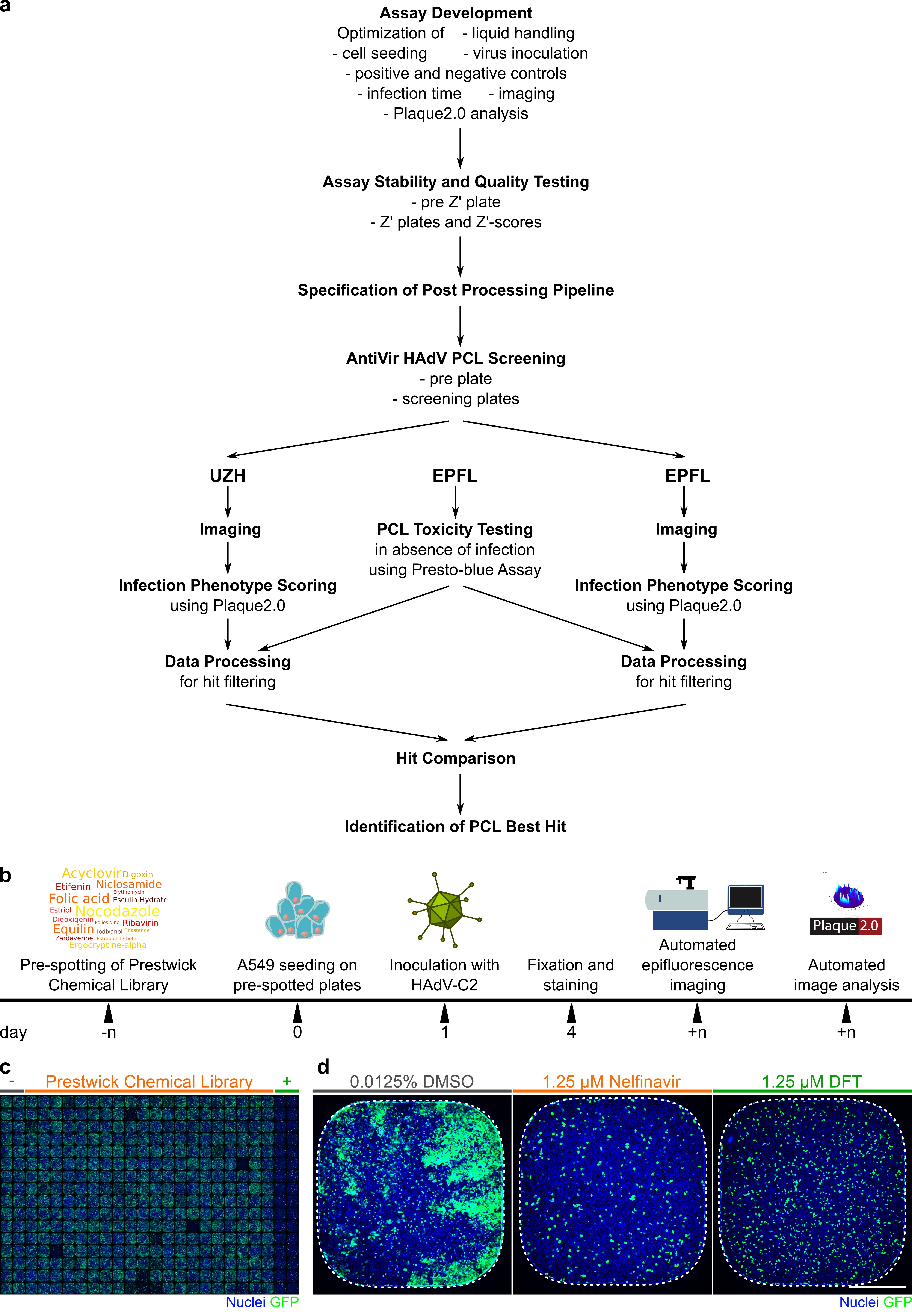

### Figure 2

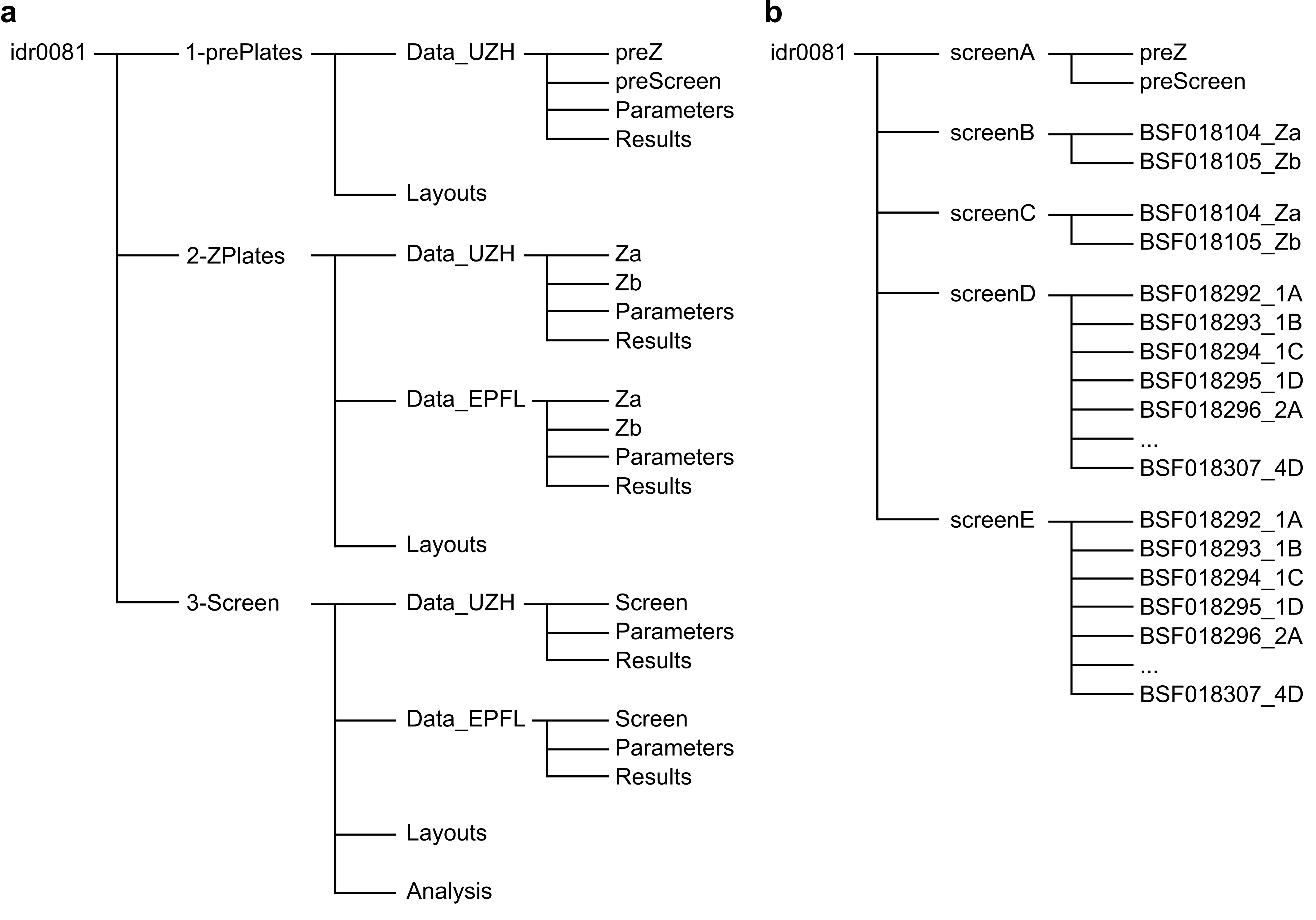

### Figure 3

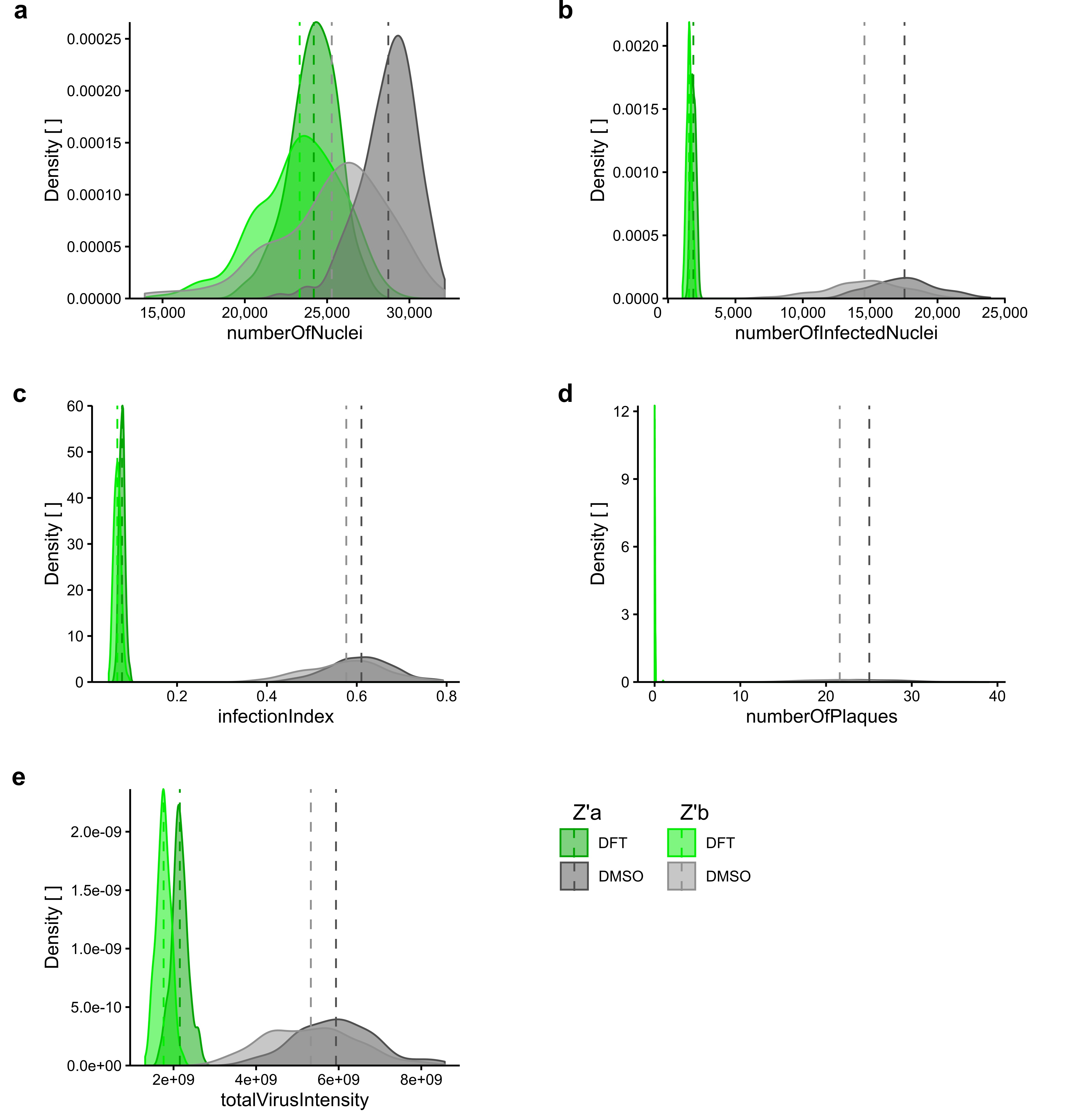

### Figure 4

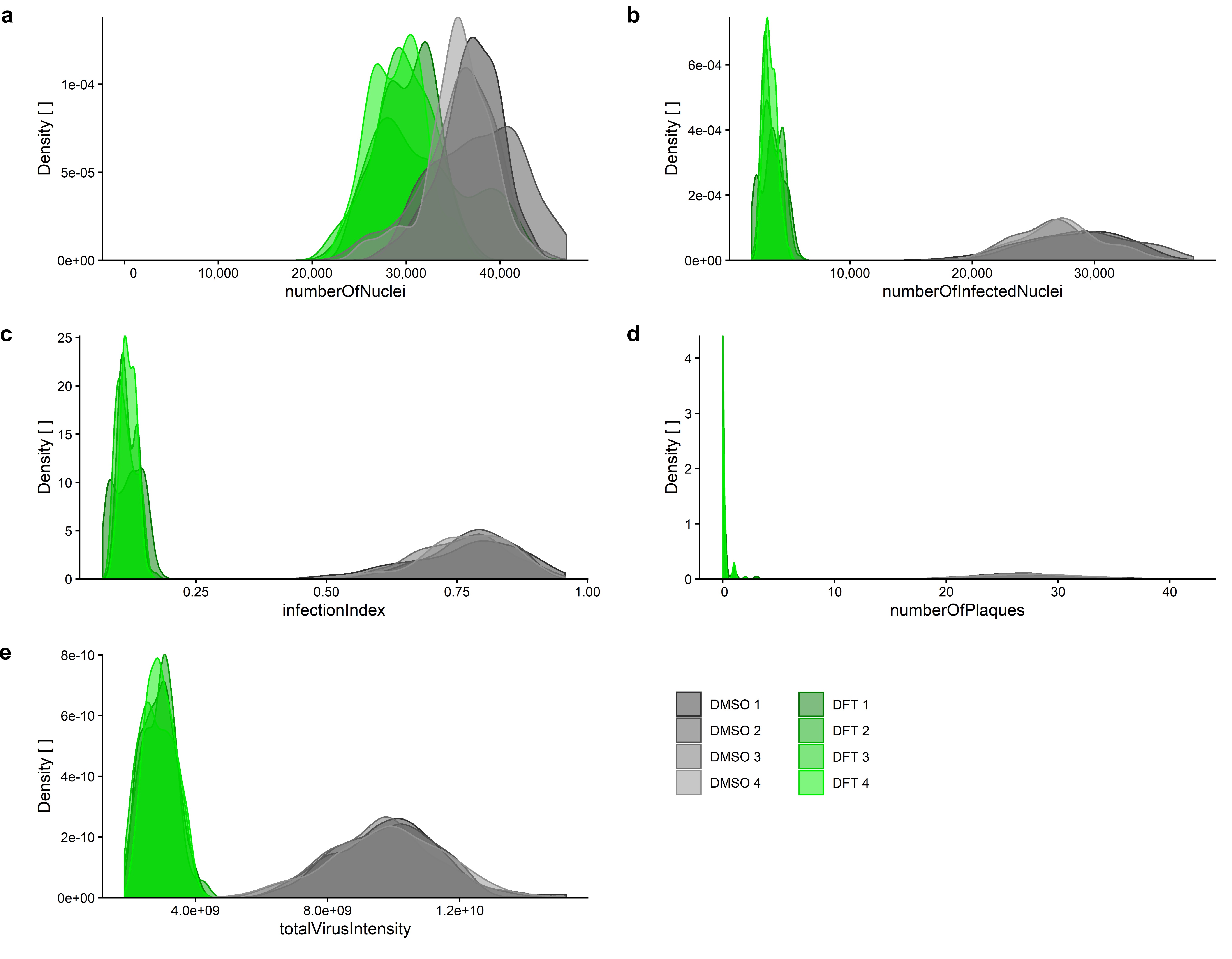

### Figure 5

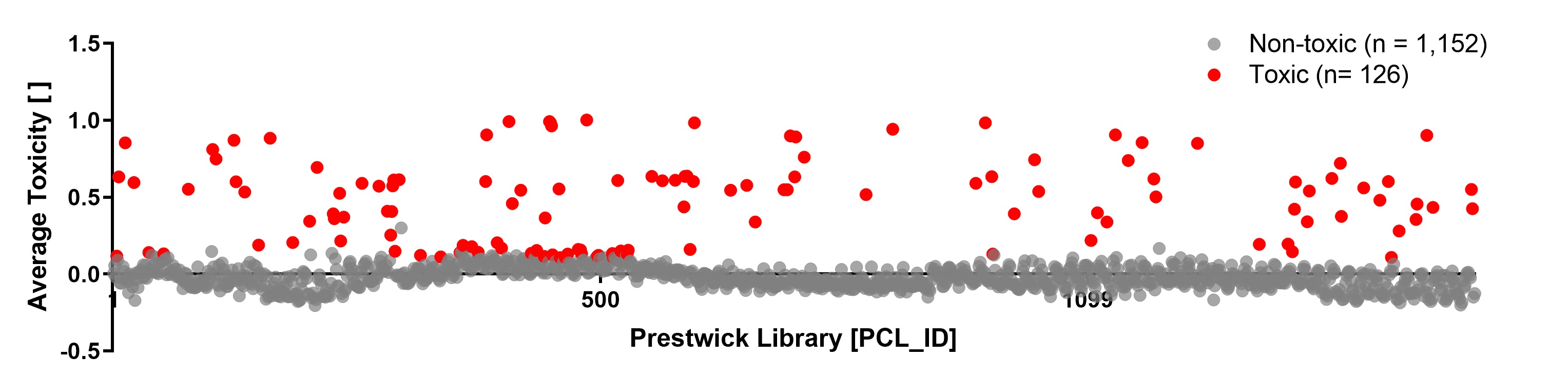

### Figure 6

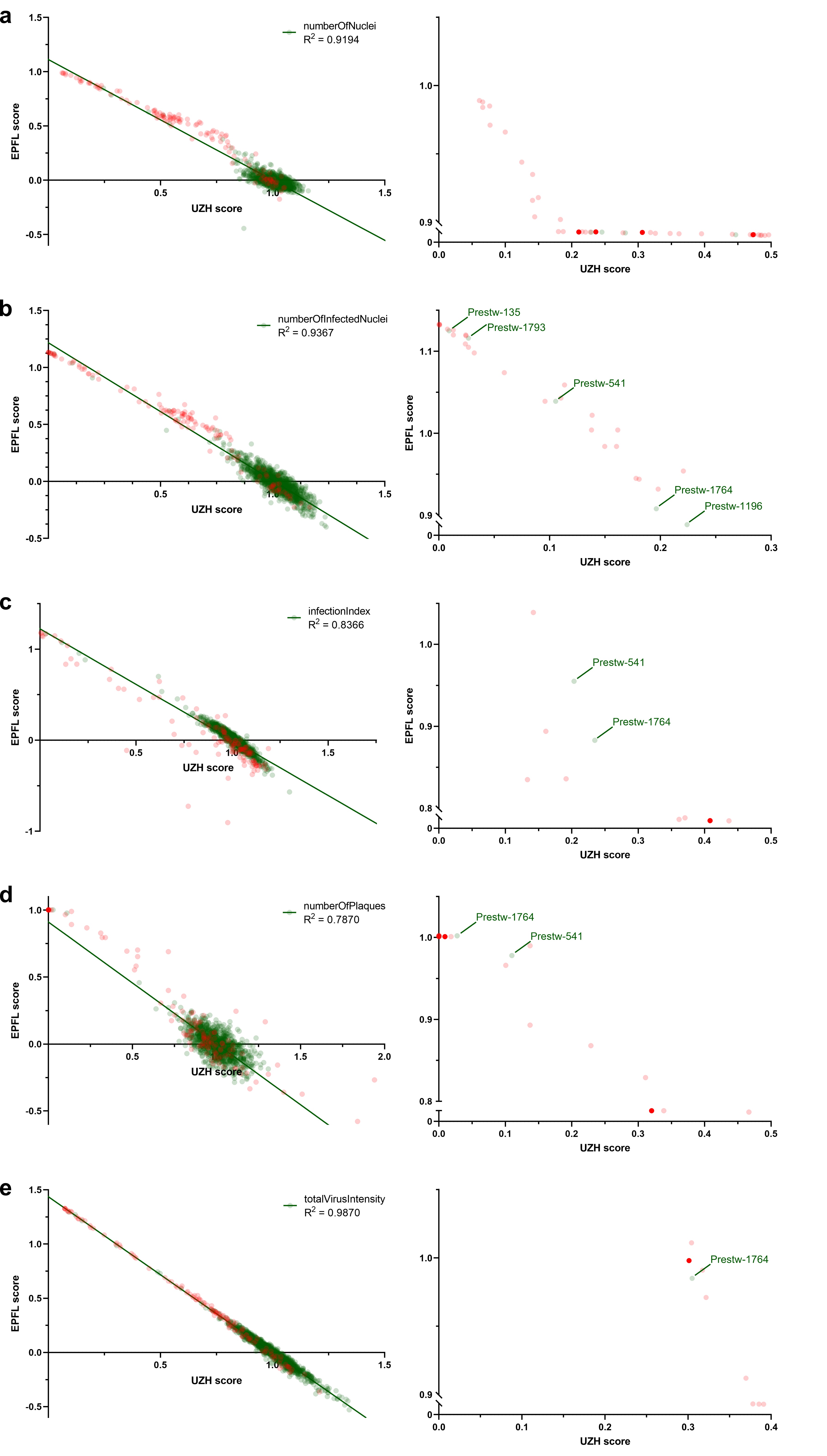
